# Sex differences in immune function and disease risk are not easily explained by an evolutionary mismatch

**DOI:** 10.1101/2024.02.13.580011

**Authors:** Audrey M. Arner, Benjamin Muhoya, Mitchell Sanchez Rozado, Michael Gurven, John Kahumbu, Joseph Kamau, Patriciah Kinyua, Francis Lotukoi, Dino J. Martins, Charles Miano, Michael J. Montague, Sospeter N. Njeru, Julie Peng, Peter Straub, Marina M. Watowich, Lea K. Davis, Noah Snyder-Mackler, Julien F. Ayroles, Amanda J. Lea

**Affiliations:** Department of Biological Sciences, Vanderbilt University, Nashville, Tennessee, USA; Evolutionary Studies Initiative, Vanderbilt University, Nashville, Tennessee, USA; Lewis Sigler Institute for Integrative Genomics, Princeton University, Princeton, New Jersey, USA; Mpala Research Centre, Nanyuki, Kenya; Department of Ecology and Evolution, Princeton University, Princeton, New Jersey, USA; Department of Microbiology and Medical Zoology, University of Puerto Rico, Rio Piedras, Puerto Rico, USA; Department of Biology, University of Washington, Seattle, WA, USA; Department of Anthropology, University of California Santa Barbara, Santa Barbara, California, USA; Institute of Primate Research, National Museums of Kenya, Nairobi, Kenya; Turkana Basin Institute, Stony Brook University, Stony Brook, New York, USA; Centre for Traditional Medicine and Drug Research, Kenya Medical Research Institute, Nairobi, Kenya; Department of Neuroscience, University of Pennsylvania, Philadelphia, PA, USA; Vanderbilt Genetics Institute, Vanderbilt University Medical Center, Nashville, Tennessee, USA; School of Life Sciences, Arizona State University, Tempe, Arizona, USA; Center for Evolution and Medicine, Arizona State University, Tempe, Arizona, USA; School for Human Evolution and Social Change, Arizona State University, Tempe, Arizona, USA; ASU-Banner Neurodegenerative Disease Research Center, Arizona State University, Tempe, Arizona, USA

## Abstract

The “Pregnancy Compensation Hypothesis” (PCH) posits that sex differences in the mammalian immune system reflect the effects of selection on female immunity during pregnancy, during which increased immunomodulation is required to reduce immune responses to the fetus, while maintaining the ability to respond to pathogens. In humans, fewer pathogens and lower parity in urban, industrialized environments has been suggested to leave female immune systems under-stimulated, creating an “evolutionary mismatch” which exacerbates sex differences in immunity and increases female autoimmune disease risk. Yet, robust tests of this mismatch hypothesis have not been conducted. Here, we first confirmed a sex-bias in autoimmune disease prevalence in a large dataset of individuals from the United Kingdom (UK Biobank). Second, we asked whether sex differences in immune function are affected by shifts toward urban, lower-parity lifestyles in a single population—the Turkana people of northwest Kenya. We found that lifestyle alters sex differences in immune cell type proportions, but not in gene expression levels. Contrary to expectations from the PCH, parity did not predict immune physiology. We then tested for generalizable trends across natural fertility mammalian populations versus urban humans (data from Turkana, NHANES, GTEx, Batwa and Bakiga, yellow baboons, and macaques). We did not find consistent relationships between lifestyle and sex-biases in immune biomarkers or between parity and the same immune outcomes. Indeed, we found that parity predicts autoimmune disease risk in the opposite direction than expected from the PCH, with higher parity associated with higher autoimmune disease risk while adjusting for relevant socioeconomic variables in the UK Biobank. Taken together, our work suggests that while sex clearly influences immune physiology and disease risk in urban settings, these effects are not easily explained by an evolutionary mismatch. Future work addressing how sex interacts with lifestyle change to generate disease is needed.

## Introduction

Biological sex influences a wide range of complex traits and disease risk [1]. For example, in post-industrial contexts, autoimmune diseases are more common in females, while non-reproductive cancers are more common in males [2–4]. In other words, females are more susceptible to diseases characterized by immune reactivity and systemic loss of self-tolerance [4], and less susceptible to diseases moderated by immune surveillance. These differences are not easily explained by sex-specific variation in reproductive hormones, environmental exposures, X-chromosome inactivation, or immune surveillance [2, 3], although recent work has nominated antigenic triggers to underly the greater prevalence of autoimmune diseases in females [5]. However, the question of what selective forces may have driven these sex differences to arise and persist remain open. In particular, it remains unclear to what degree sex differences in disease are an evolved, adaptive feature of human biology, or instead generated by some aspect of the post-industrial environments in which they have been almost exclusively studied. Understanding these evolutionary and ecological explanations for sex differences in disease and health-related traits requires expanding our focus to include understudied populations around the world, particularly in non-industrialized contexts.

One recently proposed explanation for sex differences in autoimmune disease and cancer prevalence is the Pregnancy Compensation Hypothesis (PCH), which posits that these differences are a consequence of sex-specific selection on the immune system in eutherian mammals [6]. During placentation and pregnancy, females must engage in extensive immunomodulation to both tolerate the fetus and keep the mother healthy in the face of naturally occurring pathogen threats. Mammalian female immune and reproductive physiology thus evolved in an environment characterized by high parity and high pathogen loads, but human females in post-industrial contexts often experience extremely low parity and sterile environments (as well as many other environmental features that have diverged from our evolutionary past, such as low levels of physical activity and diets high in processed foods). Thus, the PCH hypothesizes that this radical shift in reproductive ecology, pathogen exposure, and lifestyle has generated an “evolutionary mismatch” between evolved female physiology and our modern environments, which is ultimately responsible for exacerbated sex differences in human disease. It is important to clarify that the PCH applies to all placental mammals, but posits that post-industrial humans exhibit uniquely exaggerated sex differences in immunity and disease because of evolutionary mismatch.

Female human and non-human placental mammals experience dramatic changes to their immune systems during pregnancy [7–11]. Broadly, circulating immune cell composition significantly changes during pregnancy. For example, uterine lymphocyte proliferation is suppressed during pregnancy in ruminants [12], while uterine natural killer cells that regulate blood pressure and flow to the placenta increase during pregnancy in mice [13]. In humans, pregnancy-induced hormonal changes lead to increased monocytes and granulocytes and decreased lymphocytes [14, 15]. These pregnancy-associated immune changes also influence the progression of autoimmune diseases: 75% of females with rheumatoid arthritis who become pregnant show improvement of their symptoms, although these symptoms return three months postpartum [16, 17].

Also used to support the PCH, in both human and non-human placental mammals, females typically exhibit stronger innate and adaptive immune responses relative to males, who are more susceptible to many types of infection [18–21]. For example, female rats show stronger inflammatory responses in infarcts compared to males [22], and infant female piglets have increased local immune regulation relative to males [23]. In humans, female human peripheral blood mononuclear cells (PBMCs) respond more strongly to TLR7 ligands than male PBMCs [24] by inducing higher production of interferon-α (IFNα) [25]. Additionally, female humans and mice have higher basal levels of IFNα than their male counterparts [26]. These trends also hold in non-industrial humans; for example, in Tsimane forager-horticulturists, infection with hookworm is associated with diminished proinflammatory responses to viral and bacterial stimulations in females compared to males [27]. Humans also exhibit sex differences in immune composition: human males have more circulating natural killer cells and CD8+ T cells [28, 29], while females have more circulating macrophages, CD4+ T cells, basal immunoglobulin levels, B cell numbers, and C-reactive protein [18, 28, 30–34]. These examples of higher innate and adaptive immune responses in females may therefore result in a disruption in self-tolerance and development of autoimmune diseases. Autoimmune diseases in particular involve uncontrollable CD8+ and CD4+ T cell lymphocytes (both of which are sex biased) involved with both killing virus-infected cells but also cancer cells [35, 36]. However, a complete understanding of how urban environments generate the observed sex biases in autoimmune disease is still lacking.

To explain consistent sex differences in immune physiology across mammalian species, other potential evolutionary hypotheses (beyond the PCH) have been proposed. First, the immunocompetence handicap hypothesis suggests that testosterone has a dual effect in males, controlling the development and upkeep of sexual signals while also suppressing immunity [37]. However, support for this hypothesis has been equivocal, with only certain taxa having a correlation between elevated testosterone and immunomodulation [38–40]. Second, an extension of Bateman’s principle posits that, because males increase their fitness by maximizing their mating frequency whereas females increase their fitness by maximizing their lifespan, females should evolve to invest more in immunity [41]. This hypothesis also has varying levels of support across theoretical models and empirical studies [42]. Third, Metcalf & Graham propose that the immune system has evolved to deal with two main tradeoffs: sensitivity (increasing true positive responses to pathogens can be costly if it increases false positive responses to “self”) and scale of response (greater immune deployment can more effectively control pathogens but creates greater risk of damage to the host; [43]). In this hypothesis, placental females are expected to show high sensitivity and low scale of response, while males should show the opposite, with no evolutionary mismatch observed when moving from high to low fertility & pathogen contexts. Modeling suggests that these differences in trait optima may be explained by sex-specific reproductive schedules and infection risks, but these models have yet to be tested empirically [43]. Importantly, none of these three hypotheses provide explanations for the uniquely dramatic sex differences in disease etiology in post-industrial humans and suggest that sex differences would be the same across environments.

Here, we empirically test the PCH (Figure 1) in humans and nonhuman primates. To do so, we compiled data from 1) natural fertility non-human eutherian mammals (wild yellow baboons and free-living rhesus macaques), 2) subsistence-level, natural fertility, non-industrial human groups (Turkana pastoralists in Kenya, Batwa and Bakiga agriculturalists in Uganda), and 3) urban, post-industrial human populations (urban Turkana in Kenya, urban UK individuals, urban US individuals). We tested the following four predictions that follow from the PCH:

1. First, we used the UK Biobank, containing electronic health records from ∼500,000 individuals, to confirm the underlying assumption that there is a strong sex-bias in autoimmune disease and non-reproductive cancer incidence in urban, post-industrial humans. This data is more systematic, country-level data, with direct linkages to parity.
2. Second, we tested whether humans experiencing natural fertility in high-infectious burden environments (hereafter referred to as “natural fertility & infection”) show fewer sex-differences in immune physiology relative to urban, post-industrial individuals (hereafter referred to as “urban”) using data from a single population: the Turkana of Kenya. Because the Turkana population is currently transitioning from subsistence-level to urban lifestyles, individuals of the same genetic background can be found in divergent environments allowing for within-population tests complementary to the between-population analyses described below.
3. Third, we also tested for greater magnitude of sex differences in urban versus natural fertility & infection environments using data from different non-human and human populations. These analyses estimate sex differences in immune biomarkers (blood cell type counts, proportions, and gene expression levels) within each population, and then assessed whether effect sizes consistently differed between natural fertility & infection versus urban populations.
4. Finally, because the PCH posits that post-industrial declines in parity amplify sex differences in immune physiology and disease, relative to natural fertility & infection contexts, we tested for relationships between parity and immune biomarkers (in rhesus macaques, Turkana, and urban US individuals) as well as parity and disease outcomes (in the UK Biobank).

**Figure 1:**
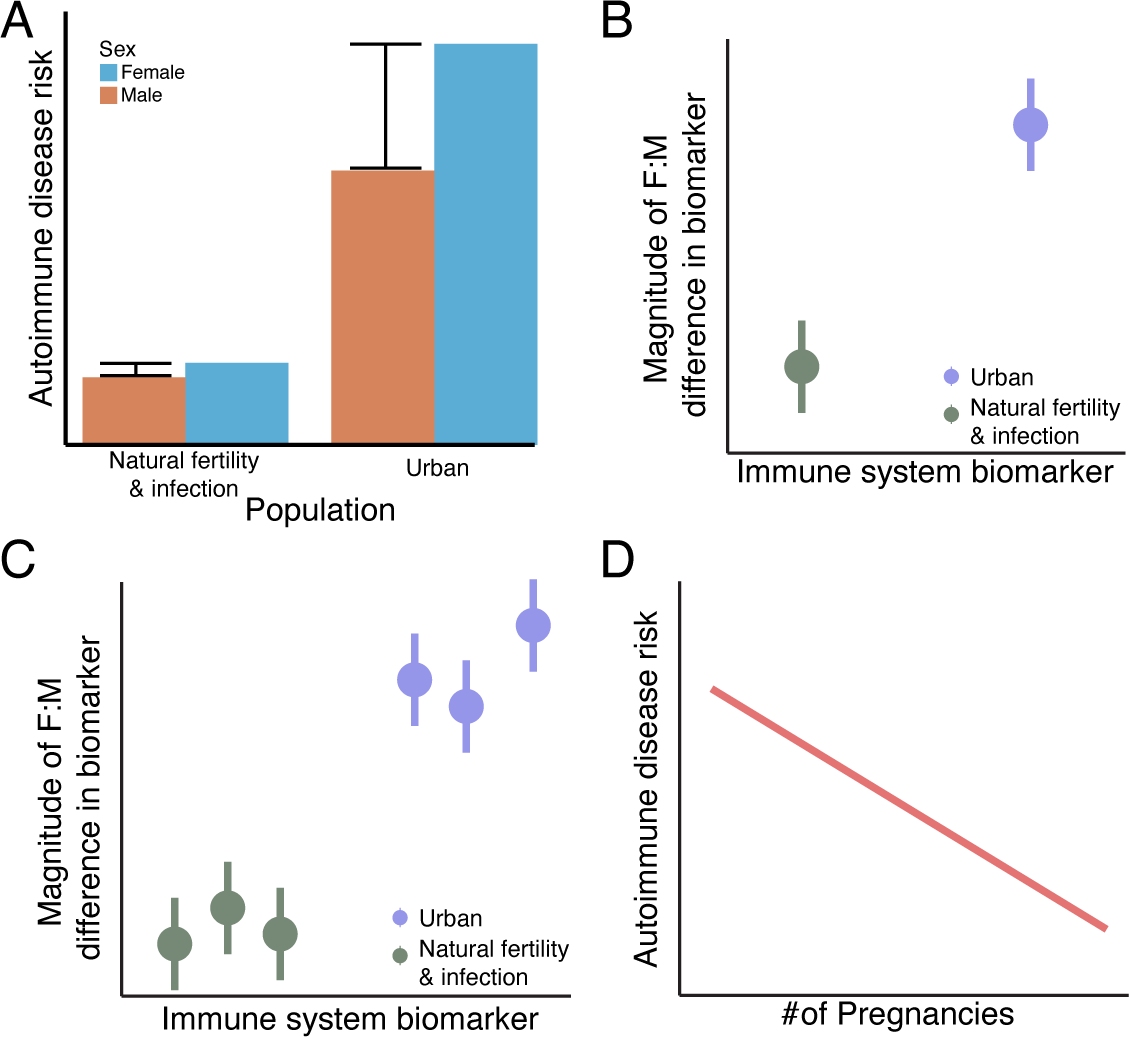
Testable predictions of the Pregnancy Compensation Hypothesis. **A.** A higher overall disease rate is expected in urban versus natural fertility & infection environments, but the magnitude of sex differences for autoimmune disease is expected to be larger in industrialized populations. **B.** Within a single population, there will be an increased magnitude of sex differences for biomarkers between groups living in urban versus natural fertility & infection environments. **C.** Across multiple genetically distinct population, the absolute value of the sex differences in biomarkers would be higher in urban versus natural fertility & infection environments. **D.** Parity will predict disease risk, such that females with more pregnancies have lower biomarkers of disease.

The PCH was first proposed by Natri et al. in 2019, but it has yet to be comprehensively tested, in part because data from both non- and post-industrial contexts are needed to robustly address its predictions. We rigorously test the PCH here for the first time using several large and integrative datasets. Together, these analyses expand our knowledge of sex as a biological variable as well as how evolutionary mismatch shapes modern day human health variation— two questions of both biomedical and evolutionary importance.

## Results

### Overview of study populations and datasets

To test the PCH, we 1) collected immune system biomarker datasets from one new human and one new non-human primate population and 2) amassed and synthesized immune system biomarker datasets from the literature for four human populations and one non-human primate population. These immune system biomarkers included data on immune cell heterogeneity (cell type proportions estimated from blood smears or flow cytometry) as well as immune cell gene expression (Figure 2A; Supplementary Table 1).

**Figure 2.**
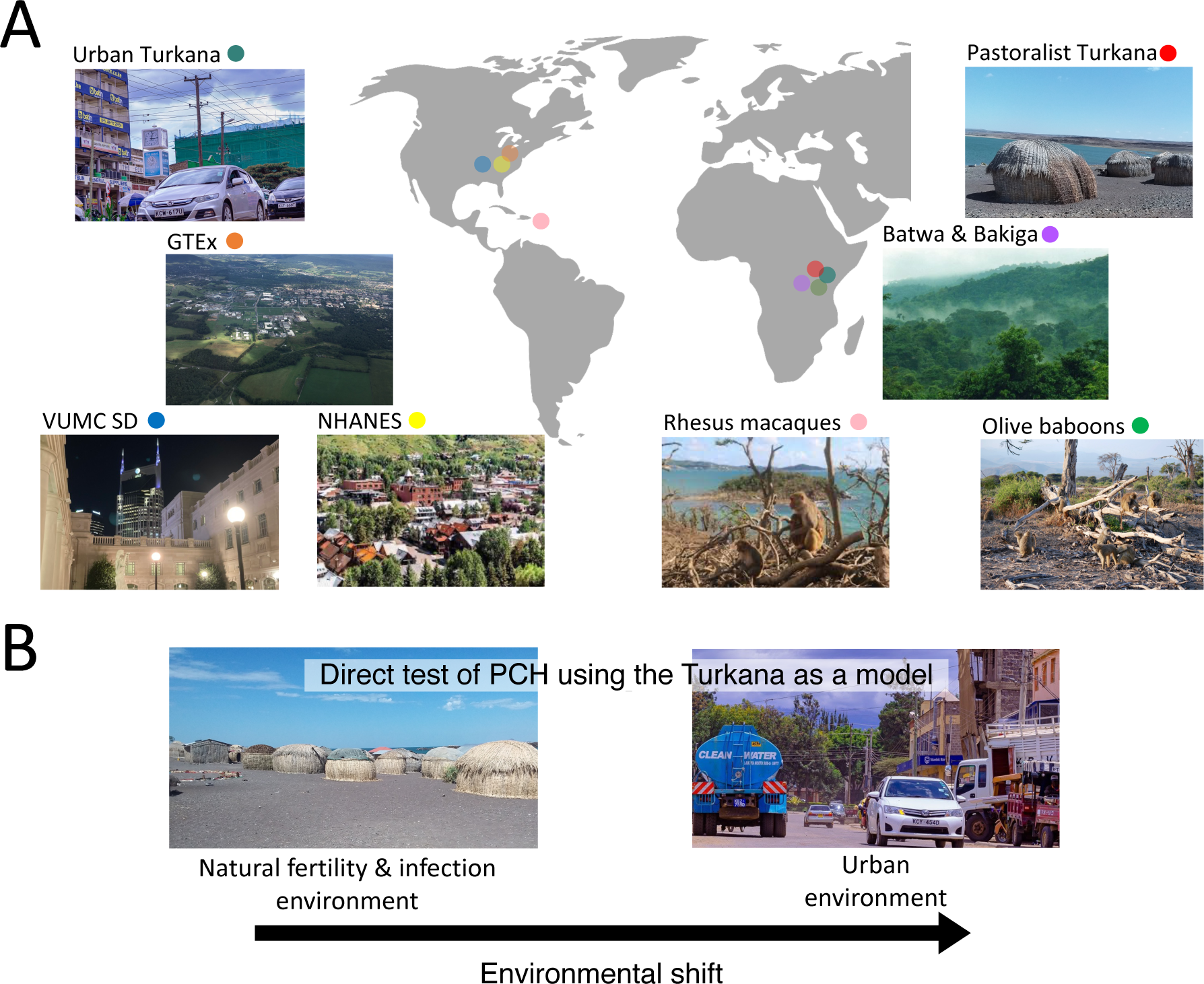
Overview of study populations and the evolutionary mismatch model. **A.** Map depicting locations of each study population. **B.** Evolutionary mismatch is currently occurring among the Turkana, such that individuals with the same genetic background live in both natural fertility & infection and urban environments

We used previously published data from five sources. First, we analyzed de-identified electronic health records for autoimmune disease, cancer, and parity data from the UK Biobank [44]. The ∼500,000 individuals included in the UK Biobank are a post-industrial cohort living in the United Kingdom with ages ranging from 40-69 and therefore towards the end of their reproductive lifespan. Second, we used cell type proportion and parity data from the National Health and Nutrition Examination Survey (NHANES). NHANES is a program run by the United States’ National Center for Health Statistics that surveys a nationally representative sample of individuals in the United States, a post-industrial, urban environment. Third, we collated cell type proportion and PBMC gene expression data from the Batwa and Bakiga, two natural fertility & infection populations living in Uganda [45]. The Batwa are traditionally a subsistence-level, rainforest hunter-gatherer population that lived in the Bwindi Impenetrable Forest of Uganda. In 1991, the Batwa were removed from their traditional homelands when the Bwindi Impenetrable National Park was created to conserve the endangered mountain gorilla. Since then, the Batwa primarily live in near settlements occupied by Bakiga subsistence-level agriculturalists. Fourth, blood cell type proportion and PBMC gene expression data were drawn from yellow baboons [46] studied by the Amboseli Baboon Research Project, who live in a natural fertility & infection environment. Finally, we used previous estimates of sex-differences in whole blood gene expression derived from the Genotype-Tissue Expression (GTEx) project [47]. GTEx is a resource of non-diseased tissues from recently deceased human donors in the US, a post-industrial, urban environment.

The first new dataset that we collected (consisting of cell type proportions, PBMC gene expression, and parity data) is from the Turkana—a subsistence-level, pastoralist population traditionally living in arid regions of northwest Kenya. Although many present-day Turkana still rely on livestock for subsistence, socioeconomic changes (e.g., the expansion of small-scale markets into northwest Kenya) have led to some Turkana relying more heavily on the market economy or even leaving the Turkana region entirely to live in more urbanized areas of central Kenya (Figure 2B). This transition provides the unique opportunity to test whether sex differences in immune physiology are amplified by urban environments, within a single population and this in the absence of confounds between genetic background and environment. Importantly, the average parity between Turkana females with completed fertility living in natural fertility & infection environments is significantly lower (n=820; beta = - 1.16; P-value = 1.01×10^-7^; linear model) than Turkana females living in urban environments (mean pregnancies for females age 45+ = 3.83 in urban environments and 4.99 in natural fertility & infection environments; Figure 4B).

We also generated new data to quantify blood cell composition, PBMC gene expression, and parity in rhesus macaques from the long-term field site of Cayo Santiago. Cayo Santiago is an island one kilometer off the coast of Puerto Rico, where rhesus macaques have been monitored since 1938. The rhesus macaques are a free-ranging, provisioned, natural fertility, and natural infection population [48].

### Female-biased differences in autoimmune disease and cancer incidence among an urban cohort

To test for sex-biases in disease incidence in post-industrial, urban environments, we used de-identified electronic health record data from the UK Biobank (n=502,364) to test for differences in the prevalence of common autoimmune diseases and non-reproductive tissue cancers (Figure 3; Supplementary Table 2). The sex biases of some autoimmune diseases we observed were similar to previous estimates, with females having higher incidence of the following autoimmune diseases: systemic lupus erythematosus, Hashimoto’s thyroiditis, Grave’s disease, multiple sclerosis, rheumatoid arthritis, primary biliary cirrhosis, and celiac disease (binomial model controlling for age, self-reported ethnicity, and socioeconomic status; 5% FDR). Unexpectedly, in this database we found that males have a significantly higher incidence the following autoimmune diseases when compared to females: myasthenia gravis, psoriasis, ulcerative colitis, type 1 diabetes, and ankylosing spondylitis. Sex biases in cancers followed more closely to those previously reported, with females having a significantly higher incidence of thyroid cancer and males having a significantly higher incidence of colorectal cancer, myeloma, lung cancer, kidney cancer, liver cancer, bladder cancer, and esophageal cancer (e.g., see summary of 13 previous studies in Table 1 of [6] and [2]); however, here all diseases were assessed in the same cohort and with the same methodology, allowing for direct effect size comparisons (similar to a study in urban Denmark [49]). No significant sex bias was found for melanoma. We repeated this analysis in the Vanderbilt University Medical Center’s (VUMC) Synthetic Derivative (SD), a database of electronic health records from ∼2.9 million individuals living in the metro area of Nashville, TN, and identified even more widespread sex biases (for example, with females having a significantly higher incidence of Alzheimer’s disease, myasthenia gravis, and psoriatic arthritis compared to males) using a cohort approach (see Supplementary Methods; Supplementary Figure 1; Supplementary Table 3).

**Figure 3:**
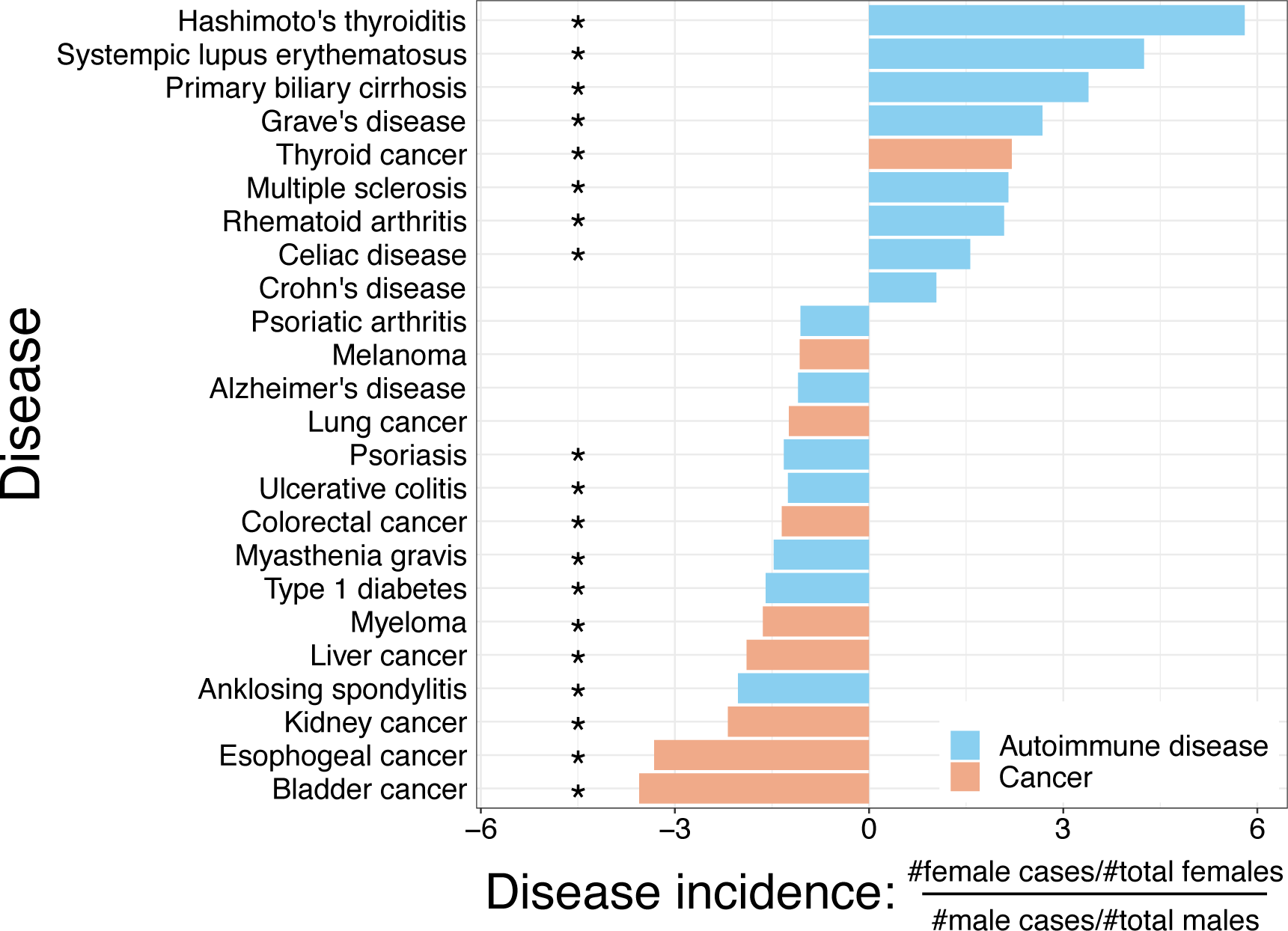
F:M ratio of disease incidence in the UK Biobank database. The given values are the ratio of the proportion of females with the disease to the proportion of males with the disease.

### Within-population effect of sex by environment interactions on immune system biomarkers

A key prediction that follows from the PCH is that sex differences in immune physiology should be more extreme in urban populations relative to populations living in environments more representative of their recent evolutionary history (e.g., natural fertility & infection). To address this prediction within a single population, we used data from the Turkana of Kenya (n=339) to test for an interaction between sex and environment (natural fertility & infection versus urban) in predicting immune biomarkers (white blood cell type proportions and PBMC gene expression data). We found that the interaction of sex and environment predicts immune physiology, such that basophil (beta = 0.3752; FDR=0.045), eosinophil (beta = 0.6029; FDR=0.001), and monocyte (beta = 0.3820; FDR=0.045) proportions exhibit distinct sex differences in natural fertility & infection versus urban settings (all in linear models controlling for age; Figure 4A). The nature of these interactions was not consistent with the PCH: sex has opposite effects on cell type proportions in the two environments, rather than consistently amplified effects in urban environments. For example, females have a lower basophil proportion in the natural fertility & infection environment when compared to males, but a higher basophil proportion in the urban environments.

When we tested for a sex x environment interaction on gene expression levels in PBMCs (n=244), we identified 6 significant genes—CDC34, ASCC2, CDKN1B, MED23, HBA2, and ID2 (FDR < 0.1; in a linear mixed-effects model controlling for relatedness, age, read count, and PC1 of cell type proportion) (Fig 4C). In contrast to what we expected under the PCH, these six genes all had greater sex biases in the natural fertility & infection environment relative to the urban environment.

**Figure 4:**
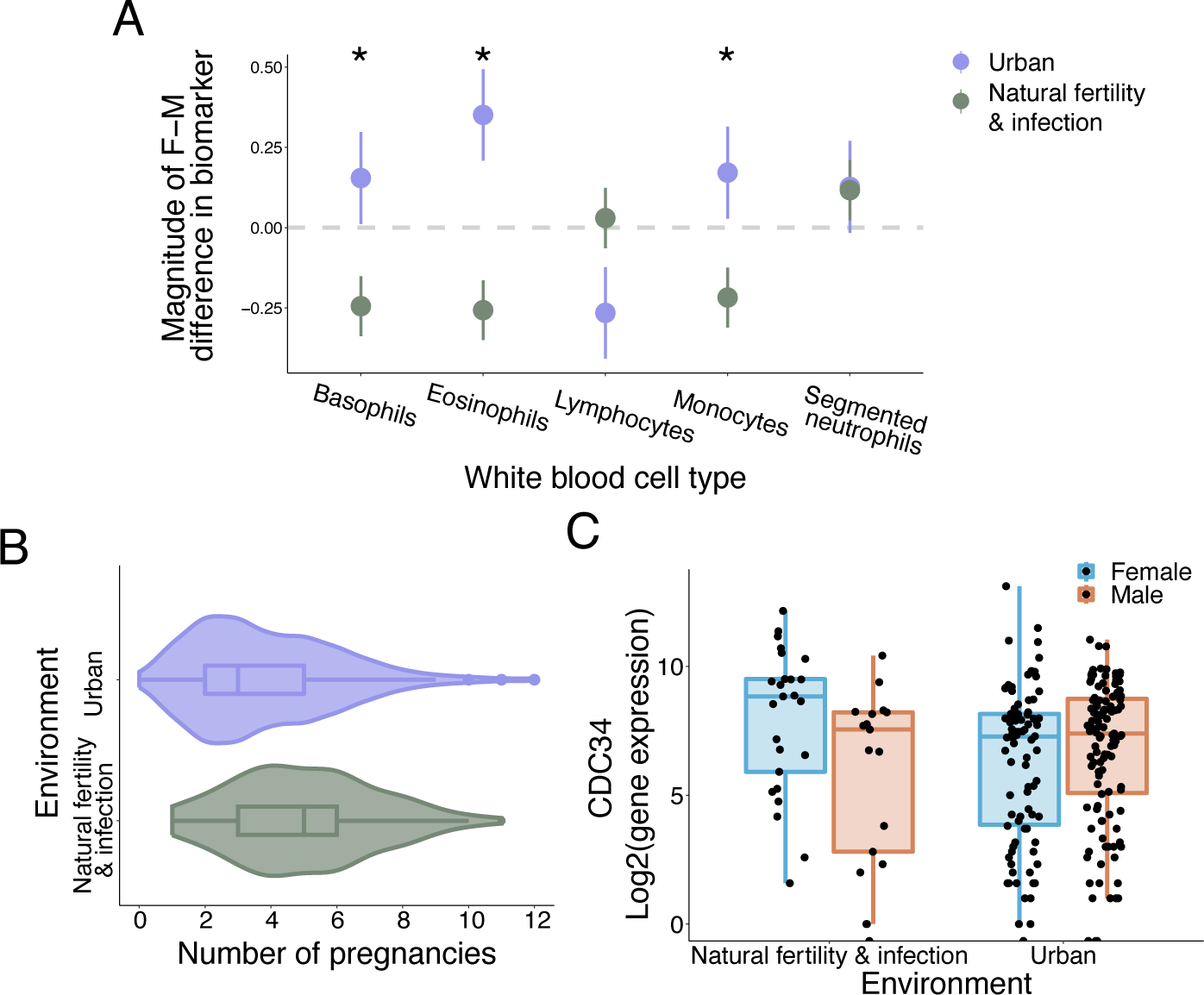
Within population sex x environment interactions on immune system biomarkers. **A.** Within white blood cell type proportions. **B.** Significant difference in number of pregnancies between urban Turkana females and natural fertility & infection Turkana females. **C.** Example of one gene, *CDC34*, which showed a significant sex x environment interaction. The natural fertility & infection population had a larger magnitude of sex difference in gene expression.

### Between population effect of sex on immune system biomarkers

To identify whether sex differences in immune system biomarkers consistently differ between populations living in natural fertility & infection versus urban environments, we analyzed white blood cell type proportion data in four additional populations: 1) the Batwa and Bakiga, 2) NHANES, 3) yellow baboons, and 4) rhesus macaques. For each cell type and each population separately, we ran linear models to identify whether sex predicts cell type proportion (controlling for covariates included in the original studies). To identify whether natural fertility & infection human (pastoralist Turkana, Batwa and Bakiga) and non-human primate (yellow baboons and rhesus macaques) populations and urban populations (urban Turkana, NHANES) followed the same patterns, we conducted a meta-analysis on the calculated effect sizes. To account for the fact that different studies focused on different sets of cell types, these analyses were aggregated by adaptive versus innate immune system cell types. Using a meta-analysis, we found that the magnitude of sex-biases in cell type proportions could be marginally predicted by environment for innate (beta = −0.1650; P-value = 0.0575; Figure 5B) but not adaptive cell types (P-value = 0.4713; Figure 5A). For innate immune cell types, populations living in natural fertility & infection environments had a lower sex bias, as predicted by the PCH.

**Figure 5:**
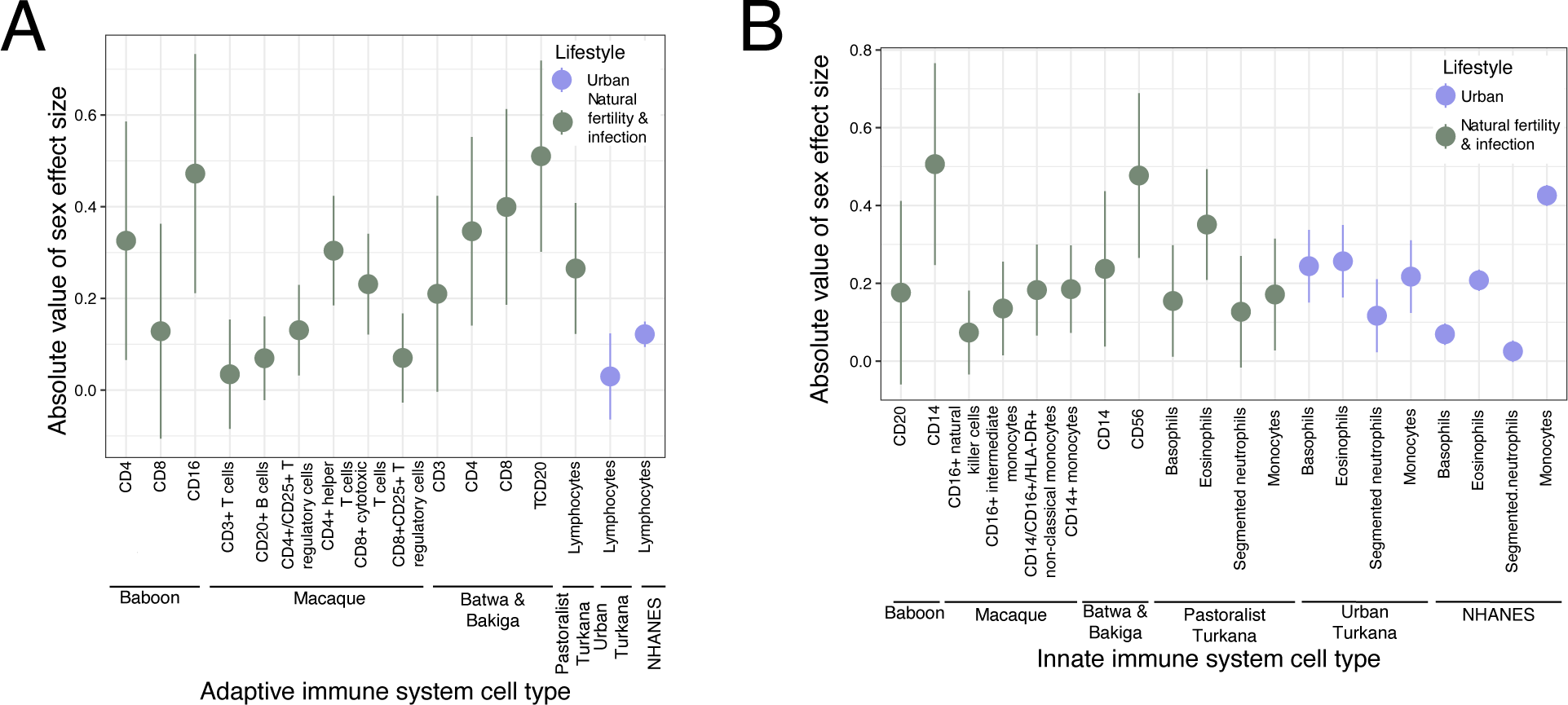
Between population analyses of sex differences in immune system biomarkers. **A.** Within adaptive immune system white blood cell count data. Cell type for each population is proportions, not absolute counts. Absolute values are shown to depict the magnitude of sex differences. **B.** Within innate immune system white blood cell count data. Cell type for each population is proportions, not absolute counts. Absolute values are shown to depict the magnitude of sex differences.

We next used a similar approach to ask whether sex-differences in immune gene expression levels consistently varied by environment across five populations: pastoralist Turkana, urban Turkana, yellow baboons, rhesus macaques, and urban individuals included in GTEx. For each population, we first ran linear models to estimate sex biases in gene expression levels for each gene. These analyses revealed minimal sex effects within each population: one gene (CDC34) was differentially expressed by sex in urban Turkana (n = 201; FDR < 0.1; in a linear mixed-effects model accounting for relatedness among individuals, age, total read count, PC1 of cell count proportion, and lane; Figure 5C), no genes were differentially expressed in natural fertility & infection Turkana (n = 43), 40 genes were differentially expressed in the Batwa and Bakiga (n=76; FDR < 0.1; in a linear model controlling for batch effects, admixture proportion, and cell type heterogeneity), no genes were differentially expressed in baboons (n=60; FDR < 0.1; in a linear mixed-effects model controlling for relatedness, age, and the first three principal components of flow data), and two genes were differentially expressed in macaques (n=176 blood samples; FDR < 0.1; in a linear model controlling for duplicate correlations, age, collection year, and sequencing depth). 1,710 sex-biased genes were previously reported for GTEx (n=948) [47].

We next conducted a meta-analysis on the sex effect sizes for genes expressed in all five populations (n=4,329 genes) and identified 266 genes where populations living in natural fertility & infection versus urban environments had consistent differences in sex-biased expression (FDR < 0.1; in a random-effects model). 55% (n=145) of these genes had stronger sex-biases in natural fertility & infection environments, while the remaining 45% (n=121) had stronger sex-biases in urban environments. Genes with stronger sex-biases in urban environments were not significantly more frequent than expected by chance (P-value = 0.16, binomial test).

To test the hypothesis that genes with environmentally dependent sex-biases would be involved in the immune system or immunomodulation, we first conducted a gene set enrichment analysis (GSEA) on our ranked list of genes (ranking on environment effect size from the meta-analysis). However, we did not find any significantly enriched pathways at a 10% FDR threshold. We next ran three distinct gene ontology (GO) analyses on 1) the 266 sex-biased genes identified in the meta-analysis, 2) the 145 genes with stronger sex biases in natural fertility & infection environments, and 3) the 122 genes with stronger sex biases in urban environments. These analyses revealed only one enriched GO term, GO:0040029: epigenetic regulation of gene expression (FDR = 0.0434), for the 122 genes with stronger sex biases in urban environments.

We next asked whether groups of genes previously identified as immune-related 1) were more likely to be significant in our meta-analyses and 2) had greater effect size estimates in our meta-analysis. We found some evidence that this was the case, using lists of genes associated with autoimmune diseases (n=470 genes) and cancers (n=96 genes) from the GWAS Catalog. Specifically, we found that genes associated with autoimmune diseases had significantly larger effect sizes for genes in our meta-analysis (P-value = 0.04125; one-sided Wilcoxon rank test; Figure 5D), as was expected via the PCH. We also found that genes associated with cancers had marginally smaller effect sizes for genes in our meta-analysis (P-value = 0.05616; one-sided Wilcoxon rank test; Figure 5E). However, we did not find a significant overlap between the 266 genes with a significant sex bias as ascertained from the meta-analysis and either autoimmune or cancer genes (P = 0.5425; P = 0.5231; Fisher’s exact test). We also tested for a significant overlap between the 121 urban sex-biased genes and either autoimmune or cancer genes. However, we again observed no significant overlap (P = 0.6841, P = 0.5231; Fisher’s exact test). These two sets of genes (those with a significant sex bias and those with a greater sex bias in the urban environment) were also not more likely to be previously identified as involved in the ex vivo response to an immune challenge (lipopolysaccharaide) (n=194 genes; P-value = 0.7756; P-value = 0.2777; Fisher’s exact tests).

### Parity effects on immune system biomarkers

To understand whether reduced parity in post-industrial environments could drive sex differences in immune physiology, as predicted by the PCH, we next tested whether immune biomarkers were indeed predicted by number of pregnancies (controlling for age; Supplementary Table 4). We found that parity does not predict any cell type proportion outcomes in Turkana females, NHANES, or rhesus macaques (all FDR > 0.1). Using a random-effects meta-analysis approach, we determined that parity predicts innate cell type proportions (beta = 0.0004; P-value = 0.0252) but not adaptive cell type proportions (beta = −0.0025; P-value = 0.3950) across populations. Using PBMC gene expression data, we did not identify any genes whose expression levels were correlated with parity in the macaques (n=81; FDR > 0.1; in a linear model controlling for duplicate correlations, age, collection year, and sequencing depth) or in the Turkana (FDR > 0.1; in a linear mixed-effects model controlling for relatedness, age, total read count, and PC1 of cell type proportion).

### Autoimmune disease incidence predicted by number of pregnancies

To directly test the prediction that low parity increases autoimmune disease risk for females in urban environments, we focused on UK Biobank females who reported number of births and stillbirths (n=273,297). We focused on the same sixteen autoimmune diseases that we tested for sex biases. Here, we found a significant effect of number of births on overall autoimmune disease risk; however, it was in the opposite direction hypothesized by the PCH, such that individuals with a greater number of pregnancies had a higher autoimmune disease risk (beta = 0.0372; P-value = 1.93×10^-4^; in a binomial model controlling for age, self-reported ethnicity, and socioeconomic status). Although this relationship was attenuated when only including autoimmune diseases with a significant sex bias in women, it was still significant (beta = 0.038; P-value = 5.87×10^-3^). To empirically test previous work that suggests fewer reproductive cycles, not necessarily due to pregnancy, attenuate autoimmune disease risk [50, 51], we then examined the relationship between autoimmune disease and age of menarche, age at menopause, total reproductive lifespan, age at first birth, age started oral contraception, and ever taken oral contraception. Of these variables, we found that lower autoimmune disease risk was associated with a younger age at first birth (beta = −0.0313; P-value = 1.32×10^-55^), younger age at menopause (beta = −0.0239; P-value = 1.02×10^-41^), and having never taken birth control (beta = −0.118; P-value = 1.03×10^-10^).

We also tested the same prediction in the VUMC SD database (n=39,452) and found a weak but significant effect such that individuals with more pregnancies were less likely to be diagnosed with an autoimmune disease (beta = −0.0022; P-value = 0.0240; in a binomial model controlling for age, self-reported race, and ethnicity). However, in this analysis we were unable to control for socioeconomic status, which is expected to be associated with a higher number of births [52] and increased autoimmune disease risk [53]. Therefore, this analysis is not fully comparable to that from UK Biobank data.

## Discussion

We used multiple immune system biomarker datasets within and between populations to provide the most comprehensive test to date of the Pregnancy Compensation Hypothesis (PCH), which posits that contemporary sex differences in autoimmune disease and cancer prevalence are a consequence of sex-specific selection on the immune system in eutherian mammals. Although our findings suggest there is potential mismatch occurring in urban environments, such that patterns of sex differences in immune physiology vary between these contexts relative to natural fertility & infection populations, our complete set of results is not fully supported by the PCH. Through our within-population analyses, we identified meaningful differences between females and males for immune system biomarkers in a single population undergoing a lifestyle transition. Specifically, we found that in the Turkana, monocyte, basophil, and eosinophil proportions exhibit distinct sex differences for natural fertility & infection versus urban individuals. However, we observed a pattern opposite of that expected under the PCH: a lower magnitude of difference in the urban relative to the natural fertility & infection environment. Additionally, although we found six genes predicted by a sex x environment interaction in the Turkana, contrary to the expectations of the PCH, all six genes had a larger sex bias in the natural fertility & infection relative to the urban population. Other potential mechanisms for explaining sex x environment effects could be hormones that interact with the immune system, differential exposure to pathogens, or behavioral, social, or cultural factors that have different sex patterns between environments and thus create sex differences in other types of exposures [54, 55]. A wide array of pathogens are widespread in the environments of most subsistence-level populations but rare in sterile post-industrial regions [56]; however, there has been little work to identify whether sex influences exposure to such pathogens.

Using between-population analyses, we found mixed evidence for the PCH. In our gene expression analyses, we found similar numbers of genes with and without a greater sex bias in urban relative to natural fertility & infection environment. However, we did not find these genes to be overrepresented among genes involved in immune physiology. Consistent with the PCH, we did find that innate immune system cell types had consistently stronger sex biases among populations in urban environments, though this effect was not particularly robust.

We found epidemiological evidence both against and in support of the predictions of the PCH. Using a large health database–the UK Biobank–we confirmed a higher incidence of most autoimmune diseases in females compared to males (and these results were confirmed in a second urban biobank, BioVU). Such findings are consistent with work evaluating sex biases in autoimmune diseases conducted across populations [2, 49, 57]. However, in the UK Biobank dataset we also found that parity predicts autoimmune disease risk in the opposite direction than expected from the PCH, with higher parity associated with higher autoimmune disease risk while adjusting for relevant socioeconomic variables. Pregnancy has been robustly shown to influence the development and expression of autoimmune diseases [58, 59], although not all autoimmune diseases are impacted the same way during and immediately following pregnancy. For example, autoimmune thyroid disease and rheumatoid arthritis have been demonstrated to improve during pregnancy but flare postpartum, while systemic lupus erythematosus worsens during pregnancy but improves postpartum [60, 61]. Previous work has also suggested that in addition to parity, longer duration of estrogen exposure, longer breastfeeding duration, and more menstrual cycles may have a protective effect against autoimmune diseases [50, 51, 62]. However, we did not find overwhelming support for this phenomenon either, with younger age of menopause, younger age at first birth, and the binary of ever taken oral contraceptive being associated with lower autoimmune disease risk. Additionally, while previous studies have found that hormonally birth control use is not associated with differences in resting levels of markers of inflammation, hormonal birth control is associated with an elevated immune response to acute stressors [63]. We note however that there are potentially important confounds, mediators, or biases not captured by the available covariates in UK Biobank, and future work is needed to untangle the precise biological links between female reproductive physiology and autoimmune disease risk, as well as to adjust for reproductive status of women in all studies.

Across our other datasets, specifically cell type proportion and gene expression data in the Turkana, rhesus macaque, and NHANES cohorts, we did not find that immune system biomarkers across lifestyles were correlated with differences in pregnancy in our observational datasets, in direct contrast with PCH predictions. This null result could be due to insufficient power to detect significant effects: while we were able to detect how parity influenced the immune system in the UK Biobank dataset (the largest cohort with 273,297 females), the other datasets we studied with parity information had much more modest sample sizes. However, we still did not detect effects when analyses were pooled in a meta-analytic framework, suggesting that if parity effects on immune physiology exist, they are extremely modest.

There are several limitations of our work. First, all datasets are observational, and all analyses are correlational, and thus cannot identify causal relationships between environment, reproductive physiology, and immune function. Future experimental work in laboratory models could be informative for testing key predictions of the PCH. Second, although the PCH posits that sex differences in autoimmune disease prevalence will be significantly lower in natural fertility & infection environments when compared to urban environments due to a mismatch in parity, we were unable to specifically analyze autoimmune prevalence in natural fertility & infection environments. These data are difficult to collect, due to a need for specialized tests and physician visits to obtain a diagnosis. Previous work has suggested lifestyle effects on autoimmune diseases broadly–though, to our knowledge, no analyses have tested if these effects differ between males and females. For example, autoimmune diseases are more common in high income countries, where birth rates are lower [64]. Additionally, autoimmune diseases are thought to be rare or absent among subsistence-level populations due to the “hygiene” or “old friends” hypothesis, which suggests that early exposure to diverse pathogens results in immune regulation that tempers inflammatory conditions [65]. Indeed, proxies for immune stimulation, such as historical disease prevalence, parasite stress, access to sanitation facilities, and access to improved-drinking water are correlated with autoimmune diseases prevalence across countries, with higher risk found in industrialized countries and urban environments [66]. Again, future work is needed to understand how these patterns break down by sex. For example, exposure to different kind of viruses in different environments could potentially induce sex-biased immune cell dynamics (i.e. in CD4+ and CD8+ T cells) that drive the higher levels of autoimmune diseases in women but higher cancer levels in men.

In summary, we found meaningful differences between sexes for immune system biomarkers quantified in individuals living in natural fertility & infection versus urban environments. While some of our results were supportive of the PCH, others were not easily interpreted in this framework. Together, our comprehensive analyses of how conserved sex differences in immune physiology are perturbed by urban, post-industrial lifestyles sheds light on our understanding of evolutionary mismatch and its consistency (or lack thereof) across populations. In other words, our results point toward heterogeneity in how lifestyle or environmental variation interacts with sex to affect physiology, which is not surprising given the diversity of ecological, behavioral, and cultural contexts across populations. Therefore, analyses that compare natural fertility & infection to urban environments are likely needed to work out population-specific biology and mechanisms, especially as they relate to how human lifestyle change impacts health. Nevertheless, our work significantly expands our knowledge of sex as a biological variable, which has clinical implications for understanding sex differences in response to immune perturbations and for answering evolutionary questions regarding disease. Future directions include research to understand the causal drivers of the sex x environment effects we observed as well as their patterning across populations. In general, our work highlights the utility and power of testing evolutionary hypotheses across cohorts, species, and data modalities, while synthesizing from the literature and newly collected data.

## Methods

### Overview of the study populations

#### Human Populations

##### UK Biobank

The UK Biobank is a large-scale, de-identified biomedical dataset containing genetic, lifestyle, and health information from ∼500,000 participants from the UK[44]. The UK Biobank originally recruited 500,000 people between ages 40-69 from 2006-2010 across the UK, during which detailed interviews were conducted about lifestyle, anthropometrics were measured, and blood, urine, and saliva samples were collected [67]. Analyses using UK Biobank data were approved by the UK Biobank (ID: 101520). Only individuals older than 18 were included in our analyses, resulting in 502,364 total people.

##### Turkana

The Turkana are a subsistence-level, pastoralist population living in a semi-arid region of northwest Kenya [68]. This environment is characterized by low annual rainfall, frequent droughts, and high temperatures year around. The Turkana are a part of the Eastern Nilotic lineages; their ancestors likely practiced nomadic pastoralism in arid regions of East Africa for thousands of years, moving into Kenya in the early 18^th^ century [68]. Present-day, traditional Turkana still rely on livestock (dromedary camels, zebu cattle, fat tailed sheep, goats, and donkeys) for subsistence and have a protein-rich diet [69]. However, as infrastructure in Kenya has improved in the past few decades, small-scale markets have expanded into northwest Kenya. These socioeconomic changes have led some Turkana to shift away from nomadic pastoralism and rely more heavily on the market economy [54]. Additionally, some individuals have left the Turkana homelands entirely and live in highly urbanized parts of central, western, and other regions of Kenya.

Interview, cell type proportion, and gene expression data were collected between April 2018 and March 2019 in Kenya’s Turkana and Laikipia counties for self-reported Turkana individuals 18 years or older. We divided individuals into three lifestyle categories as follows from previous studies of the Turkana [54]: pastoralists, urban, and rural non-pastoralists. Individuals were categorized as pastoralist if they responded in their structured interviews that they drank milk every day, their occupation was pastoralism, and they lived in Turkana County. Individuals were categorized as urban if they were sampled in Lodwar (the headquarter city of Turkana County) or lived in Laikipia County, a metropolitan area. All other individuals were designated rural non-pastoralists. We only included pastoralist (n=199) and urban (n=456) Turkana individuals in our analyses to focus on the two ends of the lifestyle spectrum. Venous whole blood samples were collected, smeared on microscope slides, fixed in methanol, and stained with Camco Stain pak. We then used a microscope to visually assess each sample for white blood cell type composition (basophils, eosinophils, lymphocytes, monocytes, and segmented neutrophils). Approximately 100 cells were counted per individual, with the total number ranging from 90-108 cells. PBMC mRNA-seq data were generated by the Turkana Health and Genomics Project (THGP) as previously described [70] (see Supplementary Methods). We had gene expression data for 43 pastoralist Turkana and 201 urban Turkana.

##### Batwa and Bakiga

The Batwa are a subsistence-level, rainforest hunter-gatherer population. Prior to 1991, the Batwa lived in the Bwindi Impenetrable Forest in Southwest Uganda for thousands of years [71]. They participated in long-term trade and inter-marrying with their Bakiga neighbors, subsistence-level agriculturalists living outside of the forest. In 1991, the Batwa were forcibly removed from the Bwindi Impenetrable National Park when it was created to conserve the endangered mountain gorilla population living inside it [72]. Since this time, the Batwa have been encouraged to live a subsistence agriculture lifestyle and live in settlements interspersed between Bakiga towns [73]. Both the Batwa and Bakiga are natural fertility populations.

Cell type composition of PBMCs were typed using conjugated antibodies, resulting in cell proportion data for the following cell types from 99 Batwa and Bakiga individuals [45]: CD3+ T cells, CD4+ T cells, CD8+ cytotoxic T cells, CD14+ monocytes, CD20+ B cells, and CD56+ natural killer cells. Harrison et al. published gene count matrices for PBMCs exposed to gardiquimod, lipopolysaccharide, and an unexposed control, and we only used the gene count matrix for the unexposed control [45]. We filtered for genes with a median log_2_(count per million) > 0.1 and removed duplicate genes and genes on the Y chromosome. These filtering steps resulted in 10,464 protein-coding genes measured in 76 individuals. We normalized read counts using the same limma-voom functions as applied to the Turkana [74]. As performed in the original study, we removed sequencing flow batch effects using ComBat and extracted the residuals for downstream modeling [75].

##### NHANES

White blood cell data was collected from the 2018 National Health and Nutrition Examination Survey (NHANES). NHANES is a program designed to assess the health and nutritional status of individuals in the United States. Beginning in 1999, the survey examines a nationally representative sample of approximately 10,000 people every two years.

We used publicly available blood cell type proportion data assessed with 5-part differential, which sorts white blood cells into subtypes using VCS technology. The white blood cell types analyzed were proportions of basophils, eosinophils, lymphocytes, monocytes, and segmented neutrophils. Only individuals older than 18 were included in our analyses, resulting in 5,263 total people.

##### GTEx

GTEx is a resource of gene expression data collected from nondiseased tissues from recently deceased human donors [81]. The project mostly contains individuals of European ancestry, but also includes up to 15% of individuals of non-European ancestry [82]. Effect sizes and standard errors for sex effects were previously published for 15,638 genes expressed in whole blood. Although whole blood and PBMCs are not the same, changes in expression profiles of both are very similar [83].

#### Non-human primates populations

##### Rhesus macaques

The rhesus macaques (*Macaca mulatta*) included in this study live on Cayo Santiago, a 15.2-ha island one kilometer off the southeastern coast of Puerto Rico. Cayo Santiago is a long-term field site where ∼1,800 free-ranging rhesus macaques have been continuously monitored between 1938-the present [76–78]. The rhesus macaques are a natural fertility population.

126 unique macaques were sampled across the two years of collection (Supplementary Methods). 50 macaques were sampled both years, resulting in a total of 176 blood samples of null condition data. We removed lowly expressed genes with fewer than 7 transcripts per million, 10 genes encoding hemoglobin and ribosomal RNA subunits, and Y-linked genes, resulting in 9,859 detectibly expressed genes for downstream analyses.

##### Yellow baboons

We analyzed blood cell proportion data and PBMC gene expression data for a longitudinally monitored population of adult yellow baboons (*Papio cynocephalus*) from the Amboseli ecosystem of southern Kenya. This population is closely related to and has some admixture with the anubis baboon (*Papio anubis*) [79]. The yellow baboons have been continuously monitored by the Amboseli Baboon Research Project (ABRP) since 1971 [80] and are a natural fertility population.

We analyzed published cell type proportions for CD4+ helper T cells, CD8+ cytotoxic T cells, CD14+ monocytes, CD16+ natural killer cells, and CD20+ B cells [46]. We also analyzed previously published PBMC gene count matrices from wild yellow baboons [46]. We only included genes that are protein-coding in *P. anubis*, genes with a median log_2_(count per million) > 0.1, and genes that were not on the Y chromosome, resulting in a total of 9,501 genes in 60 individuals.

### Sex differences in prevalence of autoimmune diseases and cancers in two cohorts

We used the UK Biobank to identify sixteen autoimmune diseases (Sjögren’s syndrome, Hashimoto’s thyroiditis, primary biliary cirrhosis, systemic lupus erythematosus, Grave’s disease, rheumatoid arthritis, multiple sclerosis, Addison’s disease, Alzheimer’s disease, celiac disease, myasthenia gravis, Crohn’s disease, Guillain–Barré syndrome, psoriasis, psoriatic arthritis, type I diabetes, ulcerative colitis, and ankylosing spondylitis) and eight cancers (thyroid cancer, colorectal cancer, lung cancer, multiple myeloma, melanoma, kidney cancer, bladder cancer, and esophageal cancer) previously identified to have sex differences across a wide range of populations [2, 6, 49]. We used a binomial regression to separately test whether each autoimmune disease or cancer could be predicted by sex (while controlling for age, self-reported ethnicity, and socioeconomic status as measured by the Townsend deprivation index.

### Within-population analysis of sex x environment interactions on white blood cell composition

We first used a within-population approach to test whether the interaction of sex and lifestyle predicts cell type proportions in the Turkana. For each of the white blood cell type proportions, we removed individuals with cell type proportions five standard deviations or more away from the mean (indicative of errors or pathology). We then scaled each cell type proportion using the “scale” function in R so that effect sizes would be standardized and comparable between white blood cell types. We used linear models to determine whether the interaction of sex and environment predict each cell type proportion separately, while controlling for age. We corrected the p-values for multiple testing using the Benjamini-Hochberg method.

### Within-population analysis of sex x environment interactions on PBMC gene expression

We next used a linear mixed-effects model that controls for genetic relatedness to identify genes significantly associated with the interaction of environment and sex using the R package EMMREML. All models corrected for age, total read count, and PC1 of cell proportion data (age and total read count were mean centered).

We checked the empirical null distribution for our analyses by permuting lifestyle ten times and rerunning our models; these analyses revealed that the genome-wide empirical null distribution of p-values was slightly non-uniform. Therefore, we used a permutation-based scheme to calculate an empirical FDR for each gene [84]. The low number of detected genes is probably somewhat but not entirely due to power, as we identified 200 genes differentially expressed by lifestyle alone (FDR < 0.1; in the same linear mixed effects model).

### Between-population analyses of sex differences in white blood cell composition

We used a between-population approach to identify whether changes in white blood cell type proportions were consistent across additional populations in natural fertility & infection and urban environments. We used a meta-analysis to examine cell type proportion data in six populations: 1) the Batwa and Bakiga, 2) NHANES, 3) yellow baboons, and 4) rhesus macaques. We split the Turkana into 5) natural fertility & infection and 6) urban individuals. For each of these white blood cell type datasets, we again removed individuals with proportions five standard deviations or more away from the mean and scaled each proportion so that effect sizes would be standardized and comparable between populations and cell types.

We tested whether sex predicts blood cell type proportions for each of our additional four populations separately. Although the specific cell types varied between populations due to differences in methodology (antibody versus flow) or an antibody panel, we conducted a meta-analysis because we were interested in the general trends observed across innate or adaptive immune system biomarkers. In our models for each population, we included the covariates that were used by the publishing research groups. Specifically, for the Batwa and Bakiga, we modeled cell type proportion as a function of sex and population (Batwa or Bakiga). Age was not included in the model, as not all Batwa and Bakiga individuals use their calendar age, and this information was thus not collected in the original study. For the NHANES and baboon datasets, we modeled cell type proportion as a function of sex and age. For macaques, we modeled cell type proportion as a function of sex, age, and data generation batch. Across datasets, we coded males as “M” and females as “F” to generate comparable effect sizes. We then conducted a mixed-effects meta-regression on the pooled set of estimated effect sizes, for adaptive and innate immune cell types separately, using the rma function in the R package *metafor* [85]. We coded the pastoralist Turkana, Batwa and Bakiga, yellow baboons, and rhesus macaques as living in a natural fertility & infection environment, while NHANES and urban Turkana were coded as living in an urban environment. We coded the following cell types as adaptive: CD3+ T cells, CD4+ helper T cells, CD8+ cytotoxic T cells, CD20+ B cells, CD4+/CD25+ T regulatory cells, CD8+/CD25+ T regulatory cells, and lymphocytes. The following cell types were coded as innate: CD14+ monocytes, CD16+ natural killer cells, CD16+ intermediate monocytes, CD14+/CD16+/HLA-DR+ non-classical monocytes, basophils, eosinophils, monocytes, and segmented neutrophils.

### Between-population analyses of sex differences in blood gene expression

We again used a between-population approach to identify whether sex effects on blood gene expression were consistent across additional populations in natural fertility & infection and urban environments. We used a meta-analysis to examine blood gene expression in four populations in addition to the two Turkana environments: 1) the Batwa and Bakiga, 2) GTEx, 3) yellow baboons, and 4) rhesus macaques. We first analyzed whether sex predicts gene expression for each population separately. In our models for each population, we included the covariates that were used by the original research groups.

Paralleling our cell type composition analyses, we split the Turkana into pastoralist (n=43) and urban (n=201) individuals to identify PBMC genes that were significantly associated with sex in these populations separately. We used linear mixed-effects models to account for relatedness, and again corrected for age, total read count, PC1, and lane, as above. We did not explore the use of an empirical FDR for these models because no differentially expressed genes were identified.

For the Batwa and Bakiga, we used linear models to test for sex effects on PBMC gene expression while controlling for fraction of reads assigned to the genome, admixture proportion, as well as CD14, CD20, and CD4 cell type proportions, which were all previously identified to impact gene expression in this dataset [45]. Prior to RNA-sequencing, cells were cultured and stimulated with medium as a control group compared to PBMCs stimulated with LPS and gardiquimod (methods in Harrison et al [45]) – we only used this null condition gene expression data. For this population we used a linear model instead of a linear mixed-effects model because we did not have access to a genomic relationship matrix. We again determined that the genome-wide distribution of p-values was slightly non-uniform by permuting the sex vector 10 times. We thus used a permutation-based scheme to calculate an empirical false discovery rate (FDR) [84].

We used previously published PBMC gene count matrices for the null condition from wild yellow baboons [46]. Following the original study, year of sample collection and sequencing depth were regressed out using the voom function in the limma package in R. We then used a linear mixed-effects model that controls for relatedness to identify genes that were significantly associated with sex, controlling for the first three principal components of the flow cytometry data. We determined that the genome-wide distribution of p-values was slightly non-uniform by permuting sex and rerunning the analysis ten times. Therefore, we used a permutation-based scheme to calculate an empirical false discovery rate (FDR) [84].

In macaques, we normalized PBMC read counts from the null condition using the same limma-voom pipeline applied to other datasets [74]. We used the duplicate correlations function in limma to control for sample identity, given that 50 individuals were sampled twice. We used a linear model, given that we did not have genetic data for these individuals, and included age, year, and sequencing depth as covariates in our linear model. We used a similar permutation approach as described above to implement an empirical FDR correction.

To include an additional urban population in our meta-analysis, we used effect sizes and standard errors of 15,638 genes estimated for GTEx whole blood RNA-seq data. The models were run with limma-voom in a pipeline comparable to our other populations. Although whole blood is not the same as PBMCs, changes in expression profiles for both tissues are similar [83].

### Meta-analyses on whether gene expression varies as a function of environment across populations

We first identified the human ortholog for each gene that was expressed in the baboon and macaque gene expression analyses. We then identified the intersection of genes that were expressed in all six of our populations, resulting in a total of 4,329 genes to be analyzed. Each population was specified to have an environment that was either natural fertility & infection (pastoralist Turkana, Batwa and Bakiga, baboon, and macaque) or urban (urban Turkana, GTEx). For each gene, we used a random-effects meta-analysis using the rma function in the R package metafor [85] to test whether the effect sizes identified in each of the previously run population-specific models could be explained by environment, followed by a Benjamini Hochberg FDR correction.

We then downloaded the GWAS catalog [86] to test whether the effect sizes from the meta-analysis were significantly different between groups of genes involved in either autoimmune or cancer disease traits previously hypothesized to have sex differences [6]. We specifically queried genes associated with the following autoimmune diseases (N=470 genes that were also present in the genes expressed among all populations): Sjogren’s syndrome, Hashimoto thyroiditis, Primary biliary cirrhosis, Systemic lupus erythematosus, Graves’ disease, Rheumatoid arthritis, Multiple sclerosis, Alzheimer’s disease, Celiac disease, Myasthenia gravis, Crohn’s disease, Psoriasis, Psoriatic arthritis, Type I diabetes, Ulcerative colitis, and Ankylosing spondylitis. The following cancers were queried (n=96 genes that were also present in the genes expressed among all populations): Thyroid cancer, Colorectal cancer, Lung cancer, Multiple myeloma, Melanoma, Kidney cancer, Bladder cancer, and Esophageal cancer. A Wilcoxon rank test was then run to test whether effect sizes from the meta-analysis differed between disease and non-disease gene sets separately for the autoimmune and cancer diseases. A Fisher’s exact test was run to test whether there is a significant overlap between genes with a significant sex bias as ascertained from the meta-analysis and autoimmune or cancer genes separately. We then ran a Fisher’s exact test to identify whether there was a significant overlap between genes with a greater sex bias in the urban environments as ascertained from the meta-analysis and autoimmune or cancer genes separately.

We next tested whether the effect sizes from the meta-analysis were significantly different between groups of genes previously identified to be differentially expressed when exposed to lipopolysaccharide (LPS) in the Batwa and Bakiga [45], baboons [46], and macaques. To be considered differentially expressed, we chose genes that had an FDR < 0.1 in all three populations. We compared against the set of genes that had an FDR > 0.1 in all three populations. When merged with our meta-analysis genes, we were left with 81 LPS-associated DE genes and 165 non-LPS-associated DE genes. We then ran a Wilcoxon rank test to identify differences in meta-analysis effect sizes between these two groups of genes. We next ran a Fisher’s exact test was run to test whether there is a significant overlap between genes with a significant sex bias as ascertained from the meta-analysis and LPS genes. Finally, we ran a Fisher’s exact test to identify whether there was a significant overlap between genes with a greater sex bias in the urban environments as ascertained from the meta-analysis and LPS genes.

We ran a GSEA to evaluate pathway enrichment of the ranked list of genes from our meta-analysis using the fgsea package in R [87]. We ran GO analyses to evaluate GO term enrichment for 1) the identified sex biased genes, 2) the genes with a greater sex bias in natural fertility & infection environments, and 3) the genes with a greater sex bias in urban environments using the clusterProfiler package in R [88].

### Analysis of parity effects on blood cell proportion across populations

To test whether age-specific fertility is correlated with markers of immune function, we first analyzed the Turkana, NHANES, and macaque white blood cell datasets. For each of these datasets, we took out males and individuals for which parity data was not recorded. The yellow baboon and Batwa and Bakiga datasets did not have available parity information, so they were not included in parity analyses. We first asked whether parity differed by lifestyle in the Turkana (n=339), while controlling for age, using a linear model. We then conducted linear models separately in each of our three populations, testing whether parity predicts white blood cell type proportions. For the Turkana data we controlled for age and lifestyle. For the NHANES data (n=2,019), we controlled only for age. For the macaques (n=81), we included both age and batch as covariates. For each population, parity and each cell count proportion was scaled so that effect sizes would be comparable. Finally, we ran a random-effects meta-analysis on the calculated effect sizes of our parity models using the rma function in the *metafor* package in R to test whether effect sizes are consistent across populations. We also ran random-effects meta-analyses on adaptive and innate cell types separately.

### Analysis of parity effects on blood cell proportion across populations

We then asked whether parity had any effects on gene expression across populations with both parity and gene expression data. In the Turkana, this included 102 females across both natural fertility & infection as well as urban lifestyles. We used a linear mixed-effects model to again account for genetic relatedness between individuals, controlling for age, lifestyle (natural fertility & infection versus urban), total read count, and PC1. Here, we used the standard Benjamini-Hochberg FDR correction, as the empirical null p-value distribution appeared uniform when permuting sex ten times. We then analyzed the effects of parity on gene expression in female macaques (N=55). Gene expression was modeled as a function of number of live births with a linear model, controlling for duplicate correlations as well as the covariates of age, year, and sequencing depth.

### Autoimmune disease incidence in BioVU and UK Biobank predicted by cycling

To test for a relationship between parity and autoimmune disease risk, we used data from the UK Biobank (n=273,297 females). To do so, we again used a binomial model controlling for age, self-reported ethnicity, and socioeconomic status (via the Townsend deprivation index). For this analysis we used all sixteen autoimmune diseases we had previously analyzed for sex bias (Type I diabetes, myasthenia gravis, ankylosing spondylitis, Crohn’s disease, multiple sclerosis, celiac disease, Hashimoto’s thyroiditis, biliary cirrhosis, psoriasis, ulcerative colitis, goiter, lupus, and rheumatoid arthritis). For this analysis we used all sixteen autoimmune diseases we had previously analyzed for sex bias. We next ran separate binomial models testing whether autoimmune disease incidence could be predicted by 1) age of first period, 2) age at menopause, 3) age at first birth, and 4) if individuals had ever taken hormonal birth control. Each binomial model again corrected for age, self-reported ethnicity, and socioeconomic status. Finally, we ran binomial models for each of our sixteen autoimmune diseases separately for each predictor to identify which diseases contributed most to negative correlations between cycling and autoimmune disease risk.

## Computational resources

Analyses were conducted in R (version 4.2.1) and with computational resources provided by The Advanced Computing Center for Research and Education at Vanderbilt University.

## Acknowledgements

This work was funded by internal awards from Princeton University to JFA. AJL was supported by internal awards from Vanderbilt University and the Searle Scholars Program. This work was supported by the National Institutes of Health (R01-AG060931; R01-AG060931-S1; R01-MH118203; R01-MH096875; R56-AG071023; F31-AG072787; R36-AG080081). AMA was supported by the National Science Foundation Graduate Research Fellowship program (DGE-1937963) and Vanderbilt University’s Evolutionary Studies Initiative. This research has been conducted using the UK Biobank Resource under Application Number 101520. The datasets used for the BioVU analyses described were obtained from Vanderbilt University Medical Center’s biorepository, which is supported by numerous sources: institutional funding, private agencies and federal grants. These include the National Institutes of Health-funded Shared Instrumentation grant S10RR025141; and Clinical and Translational Science Awards (CTSA) grants UL1TR002243, UL1TR000445 and UL1RR024975. We thank current and previous members of the Turkana Health and Genomics Project for their contributions. We are also grateful to the staff of Mpala Research Centre for their essential support. We thank the Caribbean Primate Research Center for their help in collecting data, especially James Higham and Lauren Brent. We also thank C. Jessica Metcalf for early discussion about the Pregnancy Compensation Hypothesis and the members of the Lea Lab for their discussion and comments on an earlier version of this manuscript. Above all else, we thank our participants and host communities for their hospitality and for their support in this project.

## Notes

### Competing Interest Statement

The authors have declared no competing interest.

