## Supplementary Information for "Sex differences in immune function and disease risk are not easily explained by an evolutionary mismatch"

Supplementary Methods

*Vanderbilt University Medical Center Synthetic Derivative (SD)*

The Vanderbilt University Medical Center (VUMC) Synthetic Derivative (SD) database is a de-identified version of the entire VUMC electronic health record (EHR), including records of ~3.2 million individuals. The individuals included in the database are an urban cohort from the Nashville Metro area spanning many self-described ancestries. The SD includes all clinical information in the EHR and uses an opt-out consent model that is stripped of personal identifiers [1]. The clinical system has been in use since the mid-1990s, with the SD available to researchers beginning in 2008 [1]. Analyses using EHR data from the SD were approved by the Vanderbilt Institutional Review Board (IRB #221486).

*Turkana PBMC data generation*

In brief, approximately 6mL of whole blood was collected in an EDTA tube, and PBMCs were isolated using the SepMate protocol (STEMCELL Technologies). After storage at -20C, samples were thawed and extracted with the Zymo Quick-RNA kit. RNA-sequencing libraries were prepared using a tagmentation-mediated 3' sequencing approach [2], and reads were generated on a single NovaSeq run (2x100bp sequencing) in two sequencing lanes. We trimmed adaptor sequences for low quality bases and adapter contamination using cutadapt [3]. Using STAR [4], trimmed reads were mapped to the human reference genome (hg38) and filtered for only one alignment per read. Read matrices for each sample were computed with htseq-count [5] and the GENCODE v25 GTF. Only mapped samples that had >250K reads generated were kept. We also removed duplicate samples from the same individual (always removing the sample that had a lower total read count) and samples that were missing required metadata. We obtained a genomic relatedness matrix on the same cohort of individuals previously derived from array data (to include as a random effect in our models), and we also controlled for fixed effects of age, the first PC of the genotype matrix, and cell type heterogeneity [6].

We focused our analyses on protein-coding genes. Therefore, we pruned our gene matrices to only genes that are annotated as protein coding in genome build GRCh38. Only genes with a median $\log_2$(count per million) > 0.1 were included in the analyses and duplicate genes were removed. We also removed genes that mapped to the Y chromosome, resulting in a total of 9,126 protein-coding genes. As a final quality control measure, we used a principal components analysis (PCA) to confirm that no samples were outliers. Filtered read counts were normalized using the voom function in the limma package in R. We then regressed out sequencing lane using the ComBat function and retained the residuals for downstream analyses [7].

*Macaque data generation*

Whole blood samples were collected from sedated macaques during the trap-and-release periods in 2019 and 2020. Samples were collected in 6mL K2 EDTA tubes (Beckton, Dickson and Company, cat #367899). Fresh blood samples were transported at 4C to the University of Puerto Rico-Medical Sciences campus where flow cytometric analysis was performed within six hours of sample collection. Antibodies consisted of an 8-panel cocktail that have previously been validated in rhesus macaques. The panel consisted of the following antibodies: CD20-PacBlue/Clone 2H7 (Biolegend), CD3-PerCP/Clone SP34-2 (BD), CD4-APC/Clone L200 (BD), CD8-Viogreen/Clone BW135/80 (Miltenyi), CD25-PE/Clone 4E3 (Miltenyi), CD14-FITC/Clone M5E2 (BD), CD16-PEVio770/Clone REA423 (Miltenyi), HLA-DR-APCVio770/Clone REA805 (Miltenyi). Phenotypic characterization of rhesus macaque peripheral blood mononuclear cells (PBMCs) was performed using multicolor flow cytometry. Aliquots of 150 µl of heparinized whole blood were incubated with a mix of the antibodies described for 30 minutes at 25C (room temperature). After incubation, red blood cells were fixed and lysed with the BD FACS fix and lyse solution (Cat #349202). Cells were washed twice using PBS containing 0.05% BSA and centrifuged at 1,700 RPM for 5 minutes. Cells were then processed in a MACSQuant Analyzer 10 flow cytometer (Miltenyi Biotec, CA). We specifically analyzed absolute cell type proportion data for 150 macaques for the following cell types: CD3+ T cells, CD16+ natural killer cells, CD14+ classical monocytes, CD16+ intermediate monocytes, CD14+/CD16+/HLA-DR+ non-classical monocytes, CD20+ B Cells, CD4+ helper T cells, CD8+ CD8 cytotoxic T cells, CD8+/CD25+ CD8 T regulatory cells, and CD4+/CD25+ CD4 T regulatory cells.

After blood samples were collected, 5.5mL of blood were transferred to a Leucosep tube (Greiner Bio-One Cat No. 163290) containing 3mL of Ficoll-Paque Plus (Cytiva Cat No. 17144003) and centrifuged at 1,000g for 10 min at room temperature. After centrifugation, the PBMC fraction was collected and transferred into a 50mL canonical tube and PBS was added until the volume reached 10mL. Cells were washed two times in PBS by centrifuging them at 500g for 10 minutes. Before the second wash, an aliquot of 10µl was taken, resuspended in trypan blue (Sigma, Cat No. T8154) and used to count the cells using a hemocytometer. After taking the cell count and washing for the second time, cells were resuspended in RMPI 1640 (Sigma, Cat No. R6504-10L) containing 10% FBS (Sigma Cat No. F2442) and Penicillin/Streptomycin (Fisher, Cat No. 15140122) at a concentration of 5 x $10^{-5}$ cells per mL in a new 50mL canonical tube; 1mL of the resuspended cells was then added to polypropylene free-standing tubes (VWR Cat No. 89004-302) with a final concentration of 5 x $10^{-5}$ cells.

RNA extractions were performed using the Zymo Quick-DNA/RNA MagBead kit (Zymo Catalog No. R2131), following the manufacturer's instructions with slight modifications to accommodate larger sample volumes. RNA sample concentration and quality was assessed on an AATI Fragment Analyzer. RNA sequencing libraries were prepared using the Lexogen QuantSeq-Pool Sample-Barcoded 3' mRNA-Seq Library Prep Kit (Lexogen Catalog No. 139.96), following manufacturer's instructions. 10ng of total RNA were used as RNA input and library pools were generated using a maximum of 8 samples per pool. Library pools were amplified with 15 PCR cycles in a thermocycler. All procedures were performed following the manufacturer's instructions. After library concentration was quantified, the libraries were combined in equimolar quantities and all fragments smaller than 225nt were removed in order to eliminate adapter sequences using a Pippin Prep. The resulting sample was sequenced on an Illumina NovaSeq S4 flowcell.

The sequenced reads were processed using a pipeline modeled off the Lexogen QuantSeq-Pool analysis pipeline (Lexogen-Tools 2022). After i5 and i7 indices were demultiplexed, *fastq* files were demultiplexed according to sample names and their respective i1 (detailed in Lexogen protocol) barcodes using *idemux* (version 0.1.6). Unique molecular identifiers (UMI) from the second read were extracted and written into the fastq sequence ID of the first read with *umi_tools extract* (version 1.1.2). The reads were then trimmed with *cutadapt* (version 3.6). To better capture the 3' untranslated region (UTR) and accurately count reads, a combined Ensembl and NCBI annotation of the rhesus macaque genome was generated using *AGAT* (version 0.8.0; agat_sp_merge_annotations.pl). Reads were then aligned to the combined annotation using *STAR* (version 2.7.9a) [4]. Read deduplication was performed using the UMI information in the read header with *umi_tools dedup*. The deduplicated BAM files were run through *featureCounts* (Subread version 2.0.0); only primary alignments were considered in counting, and multi-mapping reads were assigned for the primary alignment only.

*Sex differences in prevalence of autoimmune diseases and cancers in two cohorts*

We used the VUMC SD database to identify sixteen autoimmune diseases (Sjögren's syndrome, Hashimoto's thyroiditis, primary biliary cirrhosis, systemic lupus erythematosus, Grave's disease, rheumatoid arthritis, multiple sclerosis, Addison's disease, Alzheimer's disease, celiac disease, myasthenia gravis, Crohn's disease, Guillain–Barré syndrome, psoriasis, psoriatic arthritis, type I diabetes, ulcerative colitis, and ankylosing spondylitis) and eight cancers (thyroid cancer, colorectal cancer, lung cancer, multiple myeloma, melanoma, kidney cancer, bladder cancer, and esophageal cancer) previously identified to have sex differences across a wide range of populations [8–10]. We used Fisher's exact tests to identify whether there is a significant association between sex and disease outcome for each autoimmune and cancer disease separately. We report the sex ratio of each disease, which was obtained by calculating the ratio of the proportion of males with the disease to the proportion of females with the disease. Given that cancers and autoimmune diseases are more prevalent in older individuals, and thus differences in the age distribution of male and females included in the SD could bias our inference, we used a chi-squared test to confirm that the age structure was not statistically distinguishable between sexes. We did not perform age-based modeling given the structure of the data, with individuals being sub-divided into cohorts of 5 years.

*Autoimmune disease incidence in BioVU and UK Biobank predicted by cycling*

To test for a relationship between parity and autoimmune disease risk, we used data from the VUMC SD database. We used a binomial model controlling for age, self-reported race, and ethnicity to identify whether total number of preterm and term births significantly predicts presence or absence of autoimmune disease risk. For this analysis we used thirteen autoimmune diseases (Type I diabetes, myasthenia gravis, ankylosing spondylitis, Crohn's disease, multiple sclerosis, celiac disease, Hashimoto's thyroiditis, biliary cirrhosis, psoriasis, ulcerative colitis, goiter, lupus, and rheumatoid arthritis), all of which were identified by an ICD-10 code. We next ran a binomial model including only females who have had at least one preterm or term birth and a binomial model comparing autoimmune disease risk for women with no pregnancies versus women with at least on pregnancy. We then ran binomial models for each of our thirteen autoimmune diseases separately to identify which diseases contributed most to the negative correlation between parity and autoimmune disease risk. Finally, we also ran binomial models including autoimmune diseases that develop in adolescence (juvenile diabetes and psoriatic juvenile arthropathy).

Supplementary Text 1:

Throughout the text, we use the terms sex and sex differences even though we acknowledge that such designations are not binary. In the simplest sense, having a Y chromosome is what makes a mammal a male. However, sex is much more nuanced than this binary. For example, in intersex people, gene mutations can cause gonad or sexual anatomy development that does not seem to align with the sex chromosomes [11]. Despite sex being clearly on a spectrum, biological concepts of sex are important for research design that involves sex-related variables [12]. Indeed, some research institutions, such as the United States' National Institutes of Health, require considering sex as a variable in their funded research [13]. Gender is the roles and norms traditionally attributed to males and females and is considered a social construct [12]. Sex was designated for each individual in our populations in the following ways. Turkana and UK Biobank individuals self-reported their sex. The NHANES, VUMC SD, and Batwa and Bakiga cohorts self-reported their gender. If individuals reported they were a woman, we designated them as female. Individuals reporting their gender as man were designated as male. In the GTEx database, sex was designated based on the donor's identification of sex based on self-report, family/next of kin, or medical record abstraction. Sex was inferred for the yellow baboons and rhesus macaques. Sex, not gender, is used for non-human primates because gender is based on perception [14].

Supplementary Table 1

| Population | Environment | Location | N | WBC smear | WBC count | Gene expression | Parity |
|---|---|---|---|---|---|---|---|
| Pastoralist Turkana | Traditional | Kenya | 199 | X | | X | X |
| Urban Turkana | Novel | Kenya | 456 | X | | X | X |
| Batwa/Bakiga | Traditional | Uganda | 99 | | X | X | |
| NHANES | Novel | United States | 5,263 | X | | | X |
| GTEx | Novel | United States | 755 | | | X | |
| Baboon | Traditional | Kenya | 60 | | X | X | |
| Macaque | Traditional | Puerto Rico, U.S. | 126 | | X | X | X |
| BioVU | Novel | United States | 2,971,254 | | | | X |
| UK Biobank | Novel | United Kingdom | 502,364 | | | | X |

Supplementary Table 2

| Disease | Beta[1] | P-value[2] | FDR |
|---|---|---|---|
| Systemic lupus erythematosus | 1.464 | $4.1x10^{-52}$ | $1.4x10^{-51}$ |
| Hashimoto's thyroiditis | 1.756 | $2.5x10^{-49}$ | $7.5x10^{-49}$ |
| Grave's disease | 1.018 | $1.3x10^{-185}$ | $1.6x10^{-184}$ |
| Multiple sclerosis | 0.759 | $1.9x10^{-56}$ | $9.1x10^{-56}$ |
| Rheumatoid arthritis | 0.767 | $4.1x10^{-39}$ | $1.2x10^{-38}$ |
| Thyroid cancer | 0.788 | $2.4x10^{-22}$ | $3.8x10^{-22}$ |
| Primary biliary cirrhosis | 1.242 | $3.7x10^{-25}$ | $6.8x10^{-25}$ |
| Alzheimer's disease | 8.321 | 0.999 | 0.999 |
| Celiac disease | 0.456 | $6.1x10^{-37}$ | $1.5x10^{-36}$ |
| Psoriatic arthritis | -0.056 | 0.271 | 0.283 |
| Crohn's disease | 0.043 | 0.244 | 0.266 |
| Psoriasis | -0.265 | $1.8x10^{-24}$ | $3.1x10^{-24}$ |
| Ulcerative colitis | -0.220 | $3.0x10^{-16}$ | $4.0x10^{-16}$ |
| Myasthenia gravis | -0.355 | $2.8x10^{-4}$ | $3.4x10^{-4}$ |
| Type I Diabetes | -0.447 | $5.6x10^{-55}$ | $2.2x10^{-54}$ |
| Colorectal cancer | -0.268 | $2.0x10^{-28}$ | $4.0x10^{-28}$ |
| Melanoma | -0.053 | 0.076 | 0.087 |
| Myeloma | -0.462 | $1.5x10^{-19}$ | $2.1x10^{-19}$ |
| Lung and bronchus cancer | -0.164 | $1.7x10^{-10}$ | $2.1x10^{-10}$ |
| Ankylosing spondylitis | -0.693 | $3.0x10^{-33}$ | $6.5x10^{-33}$ |
| Kidney cancer | -0.754 | $4.3x10^{-70}$ | $2.6x10^{-69}$ |
| Liver cancer | -0.588 | $3.7x10^{-22}$ | $5.6x10^{-22}$ |
| Bladder cancer | -1.235 | $1.6x10^{-290}$ | $3.8x10^{-289}$ |
| Esophageal cancer | -1.164 | $9.0x10^{-110}$ | $7.2x10^{-109}$ |

[1]Beta from model, such that a positive beta is associated with higher incidence in females and a negative beta is associated with higher incidence in males

Supplementary Table 3

| Disease | Female disease ratio[1] | Male disease ratio[2] | $F_{ratio}$:$M_{ratio}$[3] | P-value[4] | FDR |
|---|---|---|---|---|---|
| Systemic lupus erythematosus | 0.0037 | 0.0007 | 5.286 | $2.2 \times 10^{-16}$ | $2.8 \times 10^{-16}$ |
| Hashimoto's thyroiditis | 0.0039 | 0.0009 | 4.333 | $2.2 \times 10^{-16}$ | $2.8 \times 10^{-16}$ |
| Grave's disease | 0.0033 | 0.0010 | 3.300 | $2.2 \times 10^{-16}$ | $2.8 \times 10^{-16}$ |
| Multiple sclerosis | 0.0045 | 0.0021 | 2.143 | $2.2 \times 10^{-16}$ | $2.8 \times 10^{-16}$ |
| Rheumatoid arthritis | 0.0082 | 0.0039 | 2.103 | $2.2 \times 10^{-16}$ | $2.8 \times 10^{-16}$ |
| Thyroid cancer | 0.0027 | 0.0013 | 2.077 | $2.2 \times 10^{-16}$ | $2.8 \times 10^{-16}$ |
| Primary biliary cirrhosis | 0.0006 | 0.0004 | 1.500 | $2.2 \times 10^{-16}$ | $2.8 \times 10^{-16}$ |
| Alzheimer's disease | 0.0027 | 0.0019 | 1.421 | $5.04 \times 10^{-13}$ | $5.04 \times 10^{-13}$ |
| Celiac disease | 0.0010 | 0.0008 | 1.25 | $2.2 \times 10^{-16}$ | $2.8 \times 10^{-16}$ |
| Psoriatic arthritis | 0.0012 | 0.0010 | 1.200 | 0.0010 | 0.0011 |
| Crohn's disease | 0.0035 | 0.0033 | 1.061 | 0.0852 | 0.0893 |
| Psoriasis | 0.0030 | 0.0031 | 0.968 | 0.1176 | 0.1176 |
| Ulcerative colitis | 0.0027 | 0.0030 | 0.800 | $4.6 \times 10^{-6}$ | $5.3 \times 10^{-6}$ |
| Myasthenia gravis | 0.0005 | 0.0006 | 0.833 | $2.2 \times 10^{-16}$ | $2.8 \times 10^{-16}$ |
| Type I Diabetes | 0.0112 | 0.0137 | 0.818 | $2.2 \times 10^{-16}$ | $2.8 \times 10^{-16}$ |
| Colorectal cancer | 0.0043 | 0.0064 | 0.672 | $2.2 \times 10^{-16}$ | $2.8 \times 10^{-16}$ |
| Melanoma | 0.0036 | 0.0055 | 0.655 | $2.2 \times 10^{-16}$ | $2.8 \times 10^{-16}$ |
| Myeloma | 0.0017 | 0.0027 | 0.630 | $2.2 \times 10^{-16}$ | $2.8 \times 10^{-16}$ |
| Lung and bronchus cancer | 0.0050 | 0.0083 | 0.602 | $2.2 \times 10^{-16}$ | $2.8 \times 10^{-16}$ |
| Ankylosing spondylitis | 0.0004 | 0.0008 | 0.500 | $2.2 \times 10^{-16}$ | $2.8 \times 10^{-16}$ |
| Kidney and renal pelvis cancer | 0.0021 | 0.0047 | 0.447 | $2.2 \times 10^{-16}$ | $2.8 \times 10^{-16}$ |
| Liver and intrahepatic bile duct cancer | 0.0013 | 0.0030 | 0/433 | $2.2 \times 10^{-16}$ | $2.8 \times 10^{-16}$ |
| Bladder cancer | 0.0013 | 0.0047 | 0.277 | $2.2 \times 10^{-16}$ | $2.8 \times 10^{-16}$ |
| Esophageal cancer | 0.0005 | 0.0017 | 0.294 | $2.2 \times 10^{-16}$ | $2.8 \times 10^{-16}$ |

[1]Ratio of females with the disease to total number of females [2]Ratio of males with the disease to total number of males [3]Sex ratio of each disease, which was obtained by calculating the ratio of the proportion of males with the disease to the proportion of females with the disease [4]P-value from the Fisher's exact test

Supplementary Table 4

| Population | Environment | Location | # females | Mean births |
|---|---|---|---|---|
| Pastoralist Turkana | Traditional | Kenya | 111 | 5.15 |
| Urban Turkana | Novel | Kenya | 228 | 3.77 |
| NHANES | Novel | United States | 2,019 | 3.48 |
| Macaque | Traditional | Puerto Rico, U.S. | 87 | 3.98 |
| BioVU | Novel | United States | 86,913 | 0.99 |
| UK Biobank | Novel | United Kingdom | 273,297 | 2.13 |

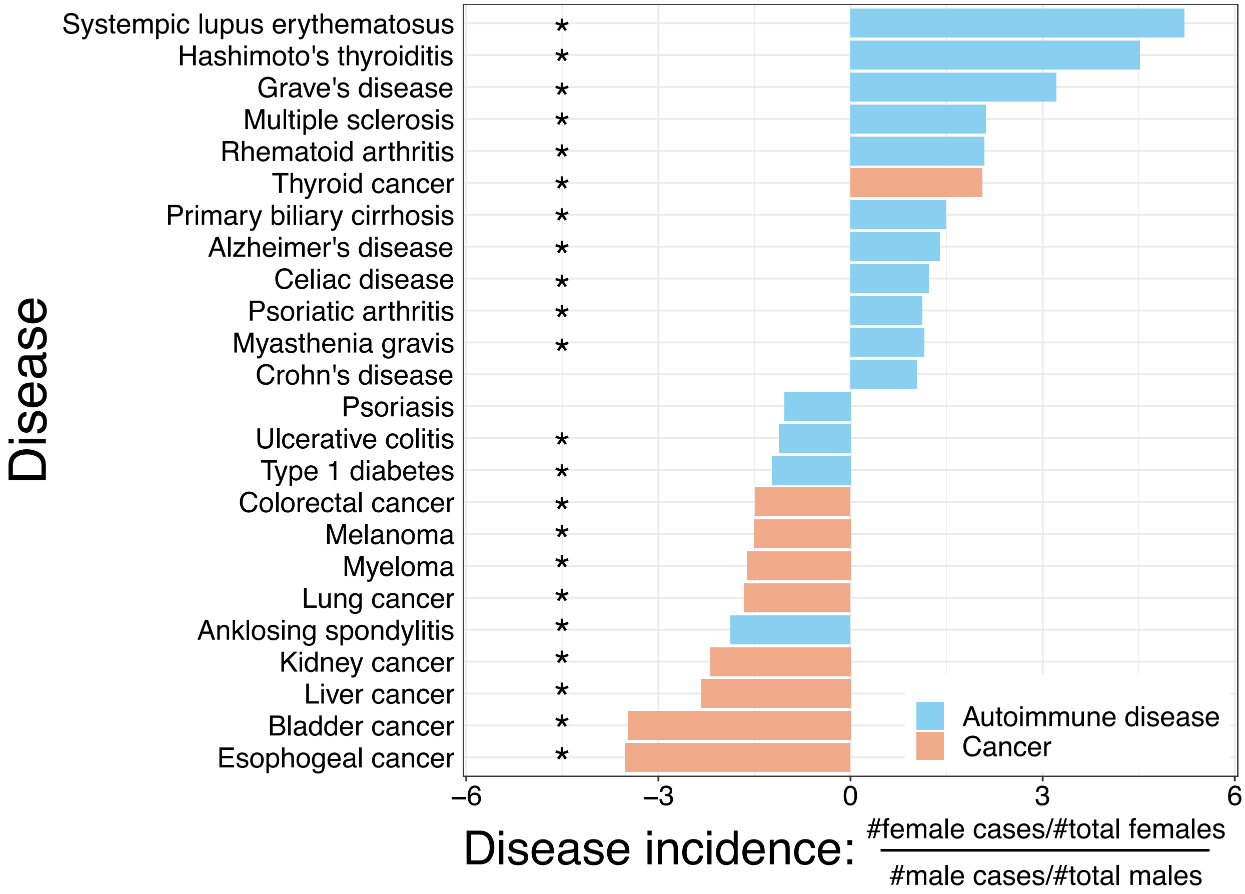


**Supplementary Figure 1: F:M ratio of disease incidence in the VUMC SD database.** The given values are the ratio of the proportion of females with the disease to the proportion of males with the disease.

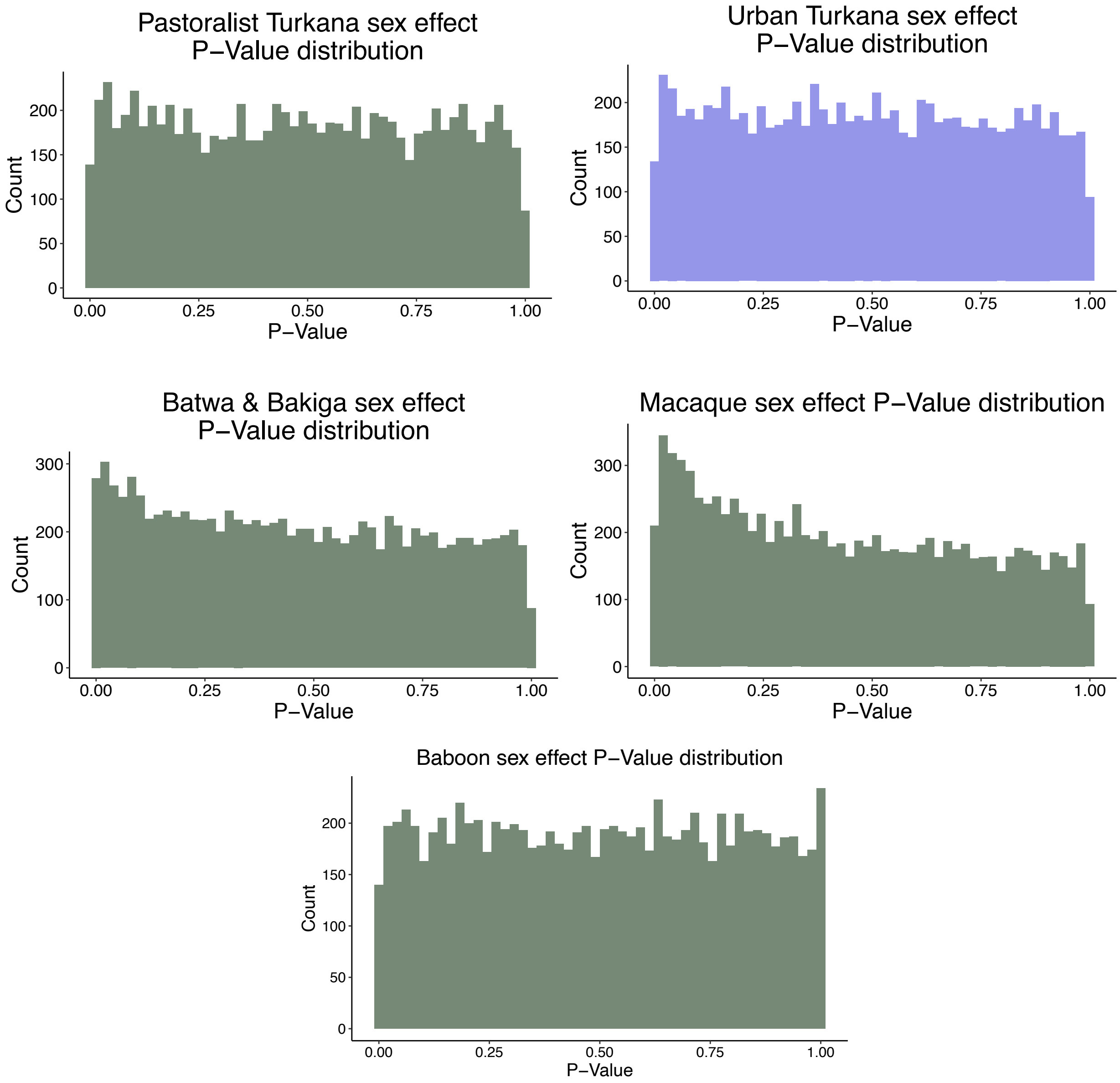


**Supplementary Figure 2: Histogram of P-Values for sex effect models from re-analyzed data.**
